# High glucose attenuates Ca^2+^ influx in cytotoxic T lymphocytes upon target recognition

**DOI:** 10.1101/2020.04.09.028019

**Authors:** Huajiao Zou, Gertrud Schwär, Renping Zhao, Dalia Alansary, Deling Yin, Eva C. Schwarz, Barbara Niemeyer, Bin Qu

## Abstract

The killing efficiency of cytotoxic T lymphocytes (CTLs) is tightly regulated by intracellular Ca^2+^ concentration. Glucose is the key energy source for CTLs, lack of which significantly impairs CTL activation, proliferation and effector functions. The impact of high glucose on Ca^2+^ influx in CTLs remains largely elusive. In this work, we stimulated primary human CD8^+^ T cells in medium containing either 25 mM (high glucose, HG) or 5.6 mM glucose (normal glucose, NG). We found that store-operated calcium entry (SOCE) induced by thapsigargin (Tg) is elevated in HG-cultured CTLs compared to their counterparts in NG. Unexpectedly, the Ca^2+^ influx elicited by recognition of target cells is reduced in HG-cultured CTLs. Under HG condition, *STIM1* and *STIM2*, the calcium sensors in the endoplasmic reticulum (ER), were down-regulated; *ORAI1*, the main structural component of calcium-release activated channels, remained unchanged, whereas *ORAI2* and *ORAI3* were up-regulated. The fraction of necrosis of HG-cultured CTLs was enhanced after killing without affecting glucose uptake. Thus, our findings reveal that HG has a distinctive impact on Tg-evoked SOCE and target recognition-induced Ca^2+^ influx in CTLs and causes more CTL death after killing, suggesting a novel regulatory role of high glucose on modulating CTL functions.

## Introduction

CD8^+^ cytotoxic T lymphocytes (CTLs) are the key players to eliminate pathogen-infected or tumorigenic cells [1, 2]. Upon recognition of cognate antigens, T-cell receptors (TCRs) are activated to trigger down-stream signaling pathways to exert their killing function. The killing processes are highly dependent on calcium [3]. At the resting state, Ca^2+^ is stored in the endoplasmic reticulum (ER) [4]. Engagement of TCRs activates phospholipase C-γ to produce inositol triphosphate (IP3) and release Ca^2+^ from the ER to the cytoplasm [5]. Store depletion activates ER Ca^2+^ sensor stromal interaction molecules (STIM) to trap and activate ORAI channels on the plasma membrane (reviewed in [6]), which let the extracellular Ca^2+^ enter the cells [6]. This depletion of Ca^2+^ ER store induced Ca^2+^ influx is termed store-operated calcium entry (SOCE).

Glucose is essential for activation, proliferation and effector functions of CD8^+^ T cells [7]. Compelling evidence shows that limiting accessibility to glucose leads to failure of elimination of tumor cells and clearance of viruses by CD8^+^ T cells [7, 8]. Elevated level of blood glucose is one of the leading symptoms of diabetes, which affects more than 400 million people worldwide [9]. Diabetic condition is associated with over-activation of CD8^+^ T cells and increased production of inflammatory cytokines [10]. Yet, the impact of high glucose (HG) on CTL function is not well understood.

In this work, we found that in HG-cultured CD8^+^ T cells, thapsigargin (Tg)-elicited SOCE was elevated, but Ca^2+^ influx evoked upon target cell recognition was decreased. With prolonged culture in HG, *STIM1* and *STIM2* were down-regulated; *ORAI2* and *ORAI3* were up-regulated, and *ORAI1* remained unchanged. The fraction of necrotic CTLs after killing was enhanced by HG without affecting glucose uptake. Our findings suggest a SOCE-involved regulatory mechanism of CTL function in the context of diabetes.

## Results and Discussion

### SOCE in CD8^+^ T cells is enhanced by high glucose

To investigate the impact of HG on Ca^2+^ influx in CTLs, we used primary human CD8^+^ T cells, stimulated with CD3/CD28 beads and cultured in NG- or HG-medium. Due to high glucose consumption by activated T cells, after day 2, glucose was added every day to keep the corresponding glucose concentration. To elicit SOCE, we used thapsigargin (Tg) to deplete Ca^2+^ from the ER. For Tg-induced SOCE, we found no difference between NG- and HG-condition for unstimulated CD8^+^ T cells (Fig. 1A-D) and at day 1.5 after stimulation (Fig. 1E-H). Remarkably, at day 3 and day 7 after bead-stimulation, Tg-induced SOCE was higher in HG-cultured CD8^+^ T cells compared to their counterparts in NG (Fig. 1I-P). We also noticed that in NG condition, upon bead-stimulation CD8^+^ T cells exhibited a significantly enhanced SOCE (Fig. 1Q), as reported by the others for Pan-T cells [11]. From day 3, SOCE in CTLs started to decline in NG-but not in HG-condition (Compare Fig. 1R and 1Q). These results suggest that Tg-elicited SOCE in CTLs is elevated by prolonged treatment of HG, which is in good agreement with the findings in mesangial cells [12].

**Figure 1.**
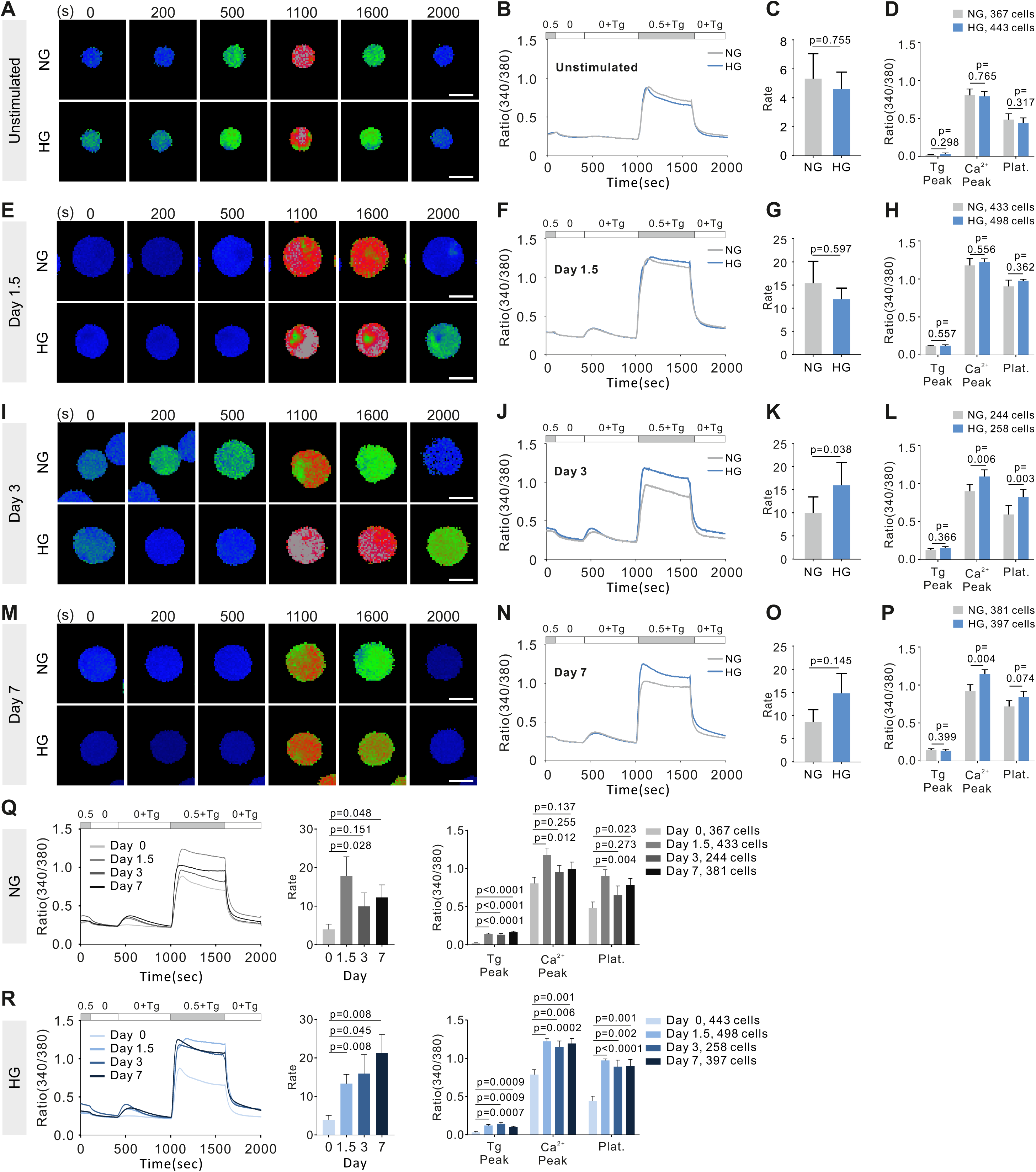
SOCE in CTLs is enhanced by HG-culture. Primary human CD8^+^ T cells were either unstimulated (**A**-**D**) or stimulated with CD3/CD28 beads in NG- (5.6 mM) or HG (25 mM)-medium for 1.5 days (**E**-**H**), 3 days (**I**-**L**) or 7 days (M-P) and then loaded with Fura-2-AM for Ca^2+^ imaging. One representative CTL (20× objective) for each condition is shown in **A, E, I** and **M**. Data were pooled from 4 independent experiments (6 donors) for **A**-**D, E**-**H** and **Q**-**R** (Day 0, Day 1.5), 5 independent experiments (7 donors) for **I**-**L** and **Q**-**R** (Day 3), and 5 independent experiments (8 donors) for **M**-**P** and **Q**-**R** (Day 7). Paired t-test was used except for **Q** and **R** (unpaired t-test). Results are presented as mean (**B, F, J, N**, and curves in **Q** and **R**) or mean ± S.E.M. (**C, D, G, H, K, L**, and bar graphs in **O** and **P**). Scale bars: 5 µm.

### Target recognition-induced Ca^2+^ influx in CTLs is reduced by high glucose

SOCE in CTLs can be also elicited by physiological stimuli such as target cell recognition. To investigated the impact of HG on target recognition-induced SOCE, we used staphylococcal enterotoxin A and B (SEA/SEB)-pulsed Raji cells as target cells [13]. We observed that at day 3 after bead-stimulation, in HG-cultured CTLs, target recognition-induced Ca^2+^ influx was significantly reduced relative to their NG-cultured counterparts, despite of the same basal level of intracellular Ca^2+^ (Fig. 2A-D). At 7 days, upon target cell engagement, we found a moderate difference in the Ca^2+^ influx rate and the Ca^2+^ peak between NG and HG-cultured CTLs, whereas no difference in Ca^2+^ plateau (Fig. 2E-H). We further confirmed that this Ca^2+^ influx was induced by target recognition since the Raji cells without SEA/SEB-pulsing did not elicit Ca^2+^ influx in CTLs (Fig. 2I, J). Together, we conclude that in CTLs, target recognition-induced Ca^2+^ influx is reduced by HG-culture.

**Figure 2.**
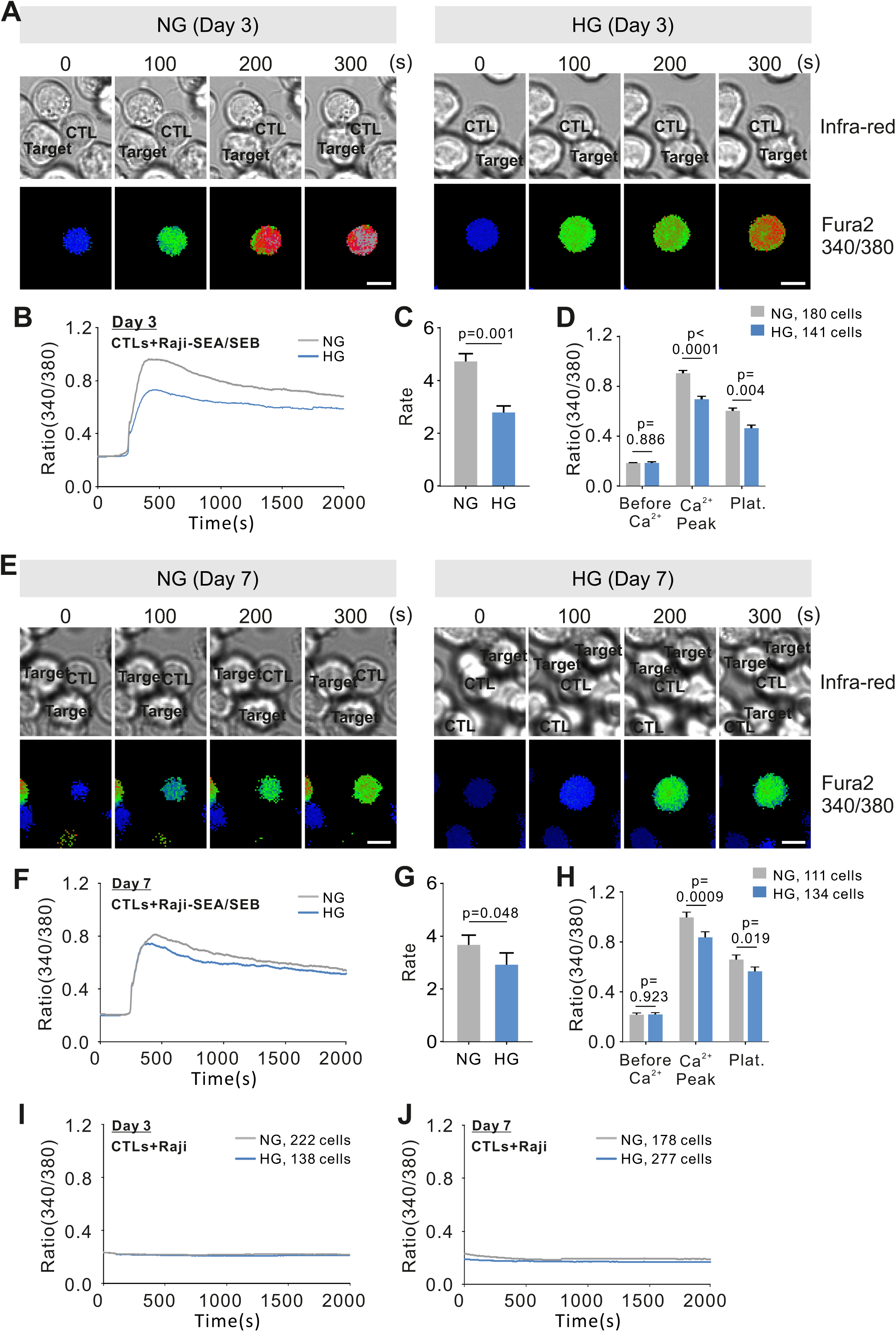
Ca^2+^ influx in CTLs upon target recognition is decreased by HG-culture. CTLs were stimulated with CD3/CD28 beads for indicated period of time in NG- or HG-medium and then loaded with Fura-2-AM for Ca^2+^ imaging. SEA/SEB-pulsed Raji cells were used as target cells (**A**-**H**). One representative CTL (20× objective) for each condition is shown in **A** and **E**. Data was pooled from 5 and 3 independent experiments (7 and 6 donors) for **A**-**D** and **E**-**H**, respectively. Raji cells without pulsing served as the negative control (pooled from 3 independent experiments (5 and 6 donors) for **I** and **J**, respectively). Paired t-test was used. Results are presented as mean (**B, F, I** and **J**) or mean ± S.E.M. (**C, D, G** and **H**). Scale bars: 5 µm.

### Expression of STIM and ORAI is distinctively regulated by high glucose

To better understand how HG could differently regulate Tg-induced SOCE and target recognition-induced Ca^2+^ influx in CTLs, we examined the expression of STIMs and ORAIs, the key components of CRAC (Ca^2+^ release-activated Ca^2+^) channels. Using quantitative PCR, we found that in HG-cultured CTLs, the predominantly expressing ORAI isoform *ORAI1* stayed unchanged, while *ORAI2* and *ORAI3* were slightly up-regulated; for the ER Ca^2+^ sensors STIM, both *STIM1* and *STIM2* were down-regulated (Fig. 3A, B). Our findings suggest that the reduction in target recognition-induced Ca^2+^ influx in CTLs by HG could be mediated by the down-regulation of STIM1 and STIM2. It is reported that ORAI2 and ORAI3 mainly function as negative-regulators of CRAC channel-mediated Ca^2+^ entry in T cells [14, 15]. Thus, up-regulation of ORAI2 and ORAI3 by HG could also contribute to reduced target recognition-induced Ca^2+^ influx, although likely to a much lesser extend due to their limited level of expression in CTLs.

**Figure 3.**
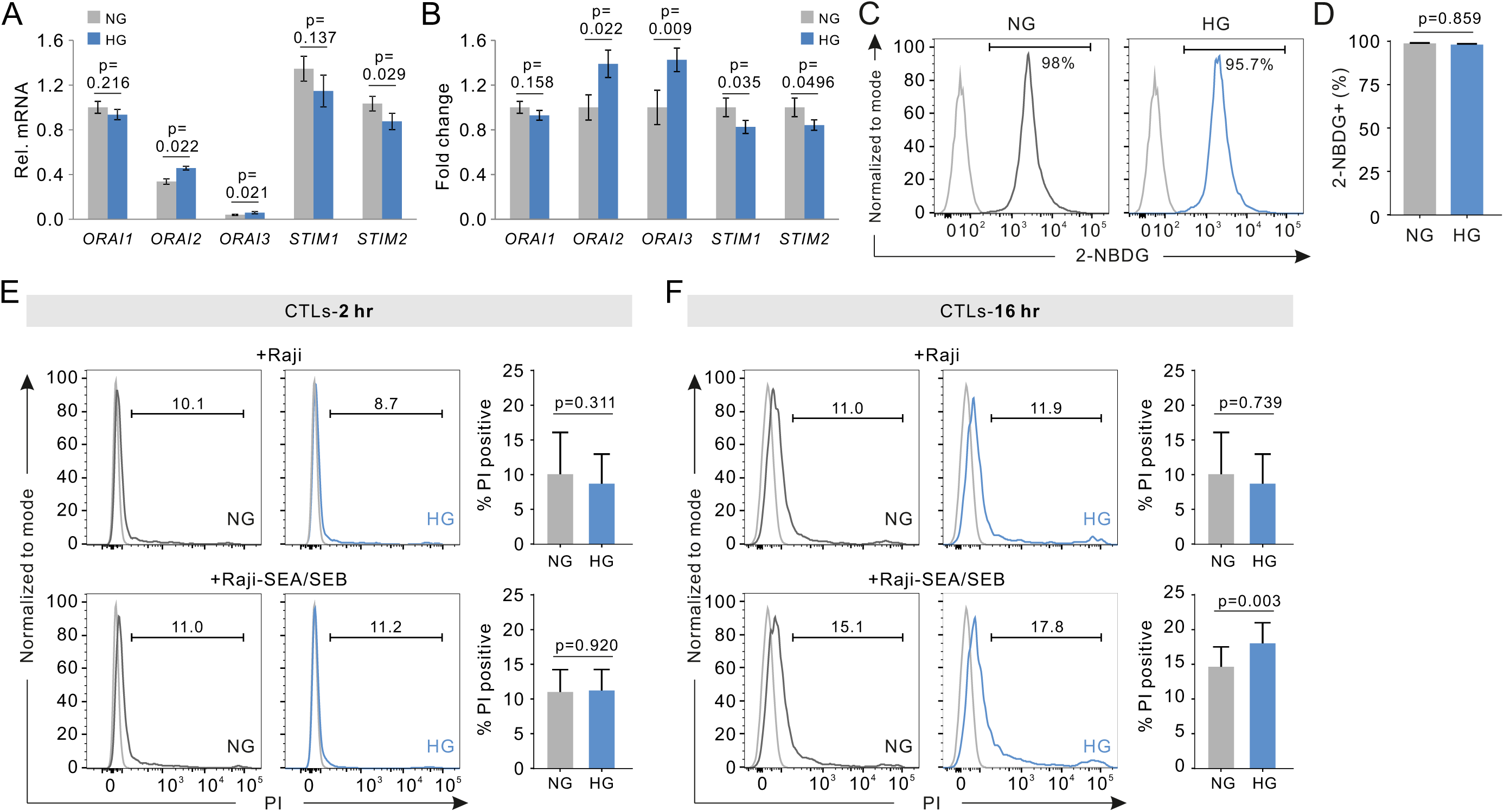
HG regulates expression of CRAC channels and impairs viability of CTLs after killing. (**A, B**) Expression of *ORAIs* and *STIMs* in CTLs. The mRNA level was determined using qPCR and either normalized to *ORAI1* in NG-cultured CTLs (**A**) or normalized to NG condition as fold change (**B**). Results are pooled from 2 independent experiments (6 donors). (**C, D**) Glucose uptake was determined by 2-NBDG. Positive cells were gated against IgG controls (gray): unstained cells. One representative donor is shown in **C.** Results are pooled from 5 independent experiments (8 donors). (**E, F**) Death of CTLs after killing. Bead-stimulated CD8^+^ T cells (CTLs) were co-incubated with indicated target cells for 2 hours (**E**) or 16 hours (**F**). PI was used to determine CTL death. Gating strategy shown in Supplementary Fig. 1. Gray: cells not stained with PI. In **E**, results are pooled from 2 and 4 independent experiments (3 and 5 donors) for Raji and Raji-SEA/SEB, respectively. In **F**, results are pooled from 3 and 5 independent experiments (4 and 6 donors) for Raji and Raji-SEA/SEB, respectively. Paired t-test was used. Results are presented as mean ± S.E.M.

Next, we sought for the functional consequence of HG-induced alteration of Ca^2+^ influx in CTLs. It is reported that ablation of SOCE by knocking out *Stim1/2* leads to severely impaired glucose uptake [16]. Using 2-NBDG, a fluorescent analog of glucose, we found that in HG and NG-cultured CTLs, glucose uptake did not differ (Fig. 3C, D), implying that the increase in SOCE by HG does not change glucose uptake.

It is reported that after glucose loading, proportion of CD8^+^ T cells is reduced in humans [17]. To examine whether HG-culture could affect viability of CTLs we used propidium iodide (PI), which binds to double-stranded DNA in dead cells. NG- or HG-cultured CTLs were co-incubated with target cells in the same medium. We found that 2 hours after killing started, not much necrosis was detected in CTLs (Fig. 3E); however, after 16 hours, the PI-positive fraction was higher in HG-cultured CTLs (Fig. 3F). Thus, our results suggest that in a time scale of several days in HG, CTLs are more prone towards necrosis, especially after execution of their killing function.

### Concluding Remarks

Collectively, in this work we report that for CTLs, HG has a distinct impact on Tg-induced SOCE and target recognition-induced Ca^2+^ influx, with the former enhanced and the latter decreased. Interestingly, under hyperglycemic conditions, calcium entry in neonatal cardiomyocytes is inhibited [18], whereas in aortic endothelial cells and mesangial cells, SOCE is enhanced [12, 19]. It indicates that the impact of HG on calcium entry varies in a cell type-dependent manner.

We found that HG-culture enhanced expression of *ORAI2* and *ORAI3* while decreased expression of *STIM1* and *STIM2*. In other cell types like aortic endothelial cells or mesangial cells, with chronic treatment of HG, STIM1-2 [12, 19] and/or ORAI1-3 [19] are upregulated. It indicates that different cell types may distinctively regulate their Ca^2+^ entry machinery in response to HG, as reflected by cell type-dependent changes in SOCE under hyperglycemic condition. The reason for enhancement of SOCE in CD8^+^ T cells by HG was not identified in this work. We postulate that splice variants of STIM could play a role. Evidence shows that down-regulation of STIM2.1, a long splice variant compared to the classic STIM2, enhances SOCE in primary human CD4^+^ T cells [20]. In addition, HG could also induce changes in Ca^2+^ signaling via superoxide anions as suggested in endothelial cells [21].

It is reported that in human umbilical vein endothelial cells, HG induces more apoptosis through SOCE [22]. Therefore, the impairment of viability of CTLs after killing by HG-culture could potentially induced through altered Ca^2+^ influx. In addition, the effect of osmolarity can be excluded since HG- and NG-cultured CTLs exhibited no difference in cell death at 2 hours. Noticeably, reduction of Ca^2+^ influx by lowering the extracellular Ca^2+^ concentration or down-regulation of ORAI1 results in an enhanced killing efficiency [3]. Whether HG could influence killing efficiency of CTLs requires further investigations.

SOCE is essential for CD8^+^ T cell activation, differentiation, proliferation and effector function [23]. Target recognition-induced or TCR activation-evoked Ca^2+^ influx is particularly important for IS formation and the consequent release of lytic granules [24]. Targeting Ca^2+^ influx is suggested as a promising strategy to boost CTL function against tumor [25]. Our findings suggest that HG may alter CD8^+^ T cell effector function and viability via regulating Ca^2+^ entry through CRAC channels, bringing new insights in understanding the regulation of CD8^+^ T cell function in the context of diabetes.

## Materials and Methods

### Antibodies and reagents

The following antibodies or reagents were used: APC/Cy7-α human CD3 (BD Bioscience), Brilliant Violet (BV) 421-α human CD19 (BioLegend), BV-421-α human CD3 (BioLegend), Fura-2-AM (Thermo Fisher Scientific), 2-NBDG (Thermo Fisher Scientific), propidium iodide (Sigma-Aldrich).

### Cell culture

Primary human CD8^+^ T cells were isolated and activated as described previously [13]. Activated CD8^+^ T cells were maintained in DMEM medium (10% FCS) with 25 mM (HG) or 5.6 mM (NG) glucose (Thermo Fisher Scientific). Raji cells were maintained in RPMI 1640 medium (10% FCS, Thermo Fisher Scientific).

### Fluorescence-based Ca^2+^ imaging

For SOCE, as described previously [15], CD8^+^ T cells were loaded with Fura-2-AM and images were taken very 5 sec for 35 min. For target recognition-induced Ca^2+^ influx [13], Fura 2-AM-loaded CTLs were settled first followed by perfusion of target cells at t = 0 sec. The images and traces were analyzed with T.I.L.L. Vision and Igor Pro6, respectively.

### Quantitative real-time PCR

Quantitative real-time PCR (qRT-PCR) was conducted as described previously [26]. As internal controls, TBP (TATA box-binding protein) and RNA polymerase were used. The mRNA level from the genes of interest is normalized to TBP and RNA polymerase as described previously [27]. The sequneces of the primers for STIM1, STIM2, ORAI1, ORAI2 and ORAI3 are described elsewhere [28].

### Flow cytometry

For glucose uptake assay, CD8^+^ T cells were loaded with 2-NBDG (120 µM) at 37°C for 30 min. For Viability assay, NG- or HG-cultured CTLs were co-incubated with target cells in AIMV medium for 2 or 16 hours. Cells were then stained with CD3 and CD19 followed by PI staining. Samples were analyzed using flow cytometry (BD FACSVerse). CD3^+^ cells were gated for further analysis using FlowJo software.

### Statistical analysis

Results were analyzed using Microsoft Excel 2016, Igor Pro6 or Prism 7 software.

## Acknowledgments

We thank the Institute for Clinical Hemostaseology and Transfusion Medicine for providing donor blood; Carmen Hässig and Susanne Renno for excellent technical help. We are grateful to Markus Hoth for constant support, inspiring discussions and advice regarding writing of the manuscript. This project was funded by the Deutsche Forschungsgemeinschaft (SFB 1027, A2 to B.Q., C4 to B.N.), Forschungsgroßgeräte (GZ: INST 256/423-1 FUGG) for the flow cytometer, and INM Fellow Program (to B.Q.).

## Ethical considerations

Research carried out for this study with human material (leukocyte reduction system chambers from human blood donors) is authorized by the local ethic committee (declaration from 16.4.2015 (84/15; Prof. Dr. Rettig-Stürmer)).

## Conflict of Interest

The authors have no financial conflicts of interest.

## Supplementary Information

## Materials and Methods

### Antibodies and reagents

All chemicals not specifically mentioned are from Sigma-Aldrich (highest grade). The following antibodies or reagents were used: APC/Cy7-conjugated anti-human CD3 antibody (BD Bioscience), Brilliant Violet (BV) 421-conjugated anti-human CD19 antibody (BioLegend), BV-421-conjugated anti-human CD3 antibody (BioLegend), Fura-2-AM (Thermo Fisher Scientific), 2-NBDG (Thermo Fisher Scientific), propidium iodide (Sigma-Aldrich).

### Cell culture

Primary human CD8^+^ T cells were prepared as described previously [1]. Briefly, primary human CD8^+^ T cells were negatively isolated from peripheral blood mononuclear cells (PBMCs) of healthy donors using Untouched Human CD8 T Cells (Miltenyi Biotec) and then activated by Dynabeads Human T-Activator CD3/CD28 (Thermo Fisher Scientific) and maintained in DMEM medium (Thermo Fisher Scientific) supplemented with 10% FBS and glucose (HG, 25 mM; NG, 5.6 mM) for 3 days if not otherwise mentioned. Raji cells were maintained in RPMI 1640 medium (Thermo Fisher Scientific) supplemented with 10% FBS and 1% penicillin–streptomycin.

### Fluorescence-based Ca^2+^ imaging

CD3/CD28 Beads were removed just before the experiments (for Day 1.5) or 24 hours prior to the experiments (for Day 3 and Day7). For SOCE measurements, as described previously [2], CD8^+^ T cells were loaded with 1 μM Fura-2-AM and then resuspended in 0.5 mM Ca^2+^ Ringer’s solution and seeded on poly-D-ornithine-coated glass coverslips for 10 min. Fluorescence emitted by 340 nm or 380 nm and infrared images were taken every 5 sec for 35 min. For target recognition-induced Ca^2+^ influx [3], Raji cells were pulsed with SEA/SEB (1 μg/ml). Fura-2-AM-loaded CD8^+^ T cells were settled on poly-D-ornithine-coated glass coverslips for 10 min followed by perfusion of Raji cells at t = 0 sec. All experiments were performed using self-built perfusion chambers with low volume and high solution exchange rate at room temperature. The captured images were analyzed by T.I.L.L. Vision software. The traces were analyzed with the software Igor Pro6.

### Quantitative real-time PCR

CTLs were harvested in TRIzol (Life Technologies) at day 3 after activation and stored at - 80°C until RNA was isolated following the manufacturer’s instructions. SuperScriptTMII Reverse Transcriptase (Life technologies) was used to generate cDNA. Subsequent quantitative PCR was conducted using the QuantiTect SYBR Green Kit (Qiagen) and a CFX96 Real-Time System (Bio-Rad). As internal controls, TBP (TATA box-binding protein) and RNA polymerase were used. The mRNA level of genes of interest is normalized to TBP and RNA polymerase and the average of normalized mRNA was taken as relative mRNA.

### Glucose uptake assay

CD8^+^ T cells were loaded with 2-NBDG (120 µM) at 37°C with 5% CO_2_ for 30 min and then washed twice with PBS containing 0.5% BSA. Then cells were incubated with BV-421 conjugated anti-CD3 antibody at 4°C for 25 min. Samples were analyzed using flow cytometry (BD FACSVerse).

### Viability assay

CD8^+^ T cells were co-incubated with Raji cells and then the cells were stained with APC/Cy7-αCD3 and BV-421-αCD19 for 25 min, followed by PI staining for 15 min at 4°C. Fluorescence intensity was detected using flow cytometry (BD FACSVerse). CD3^+^ cells were gated for further analysis using FlowJo software.

### Statistical analysis

Data are presented as the mean ± SEM if not stated otherwise. Results were analyzed using Microsoft Excel 2016, Igor Pro6 or Prism 7 (GraphPad Software, La Jolla, CA, USA) software.

---

**Supplementary Figure 1.**
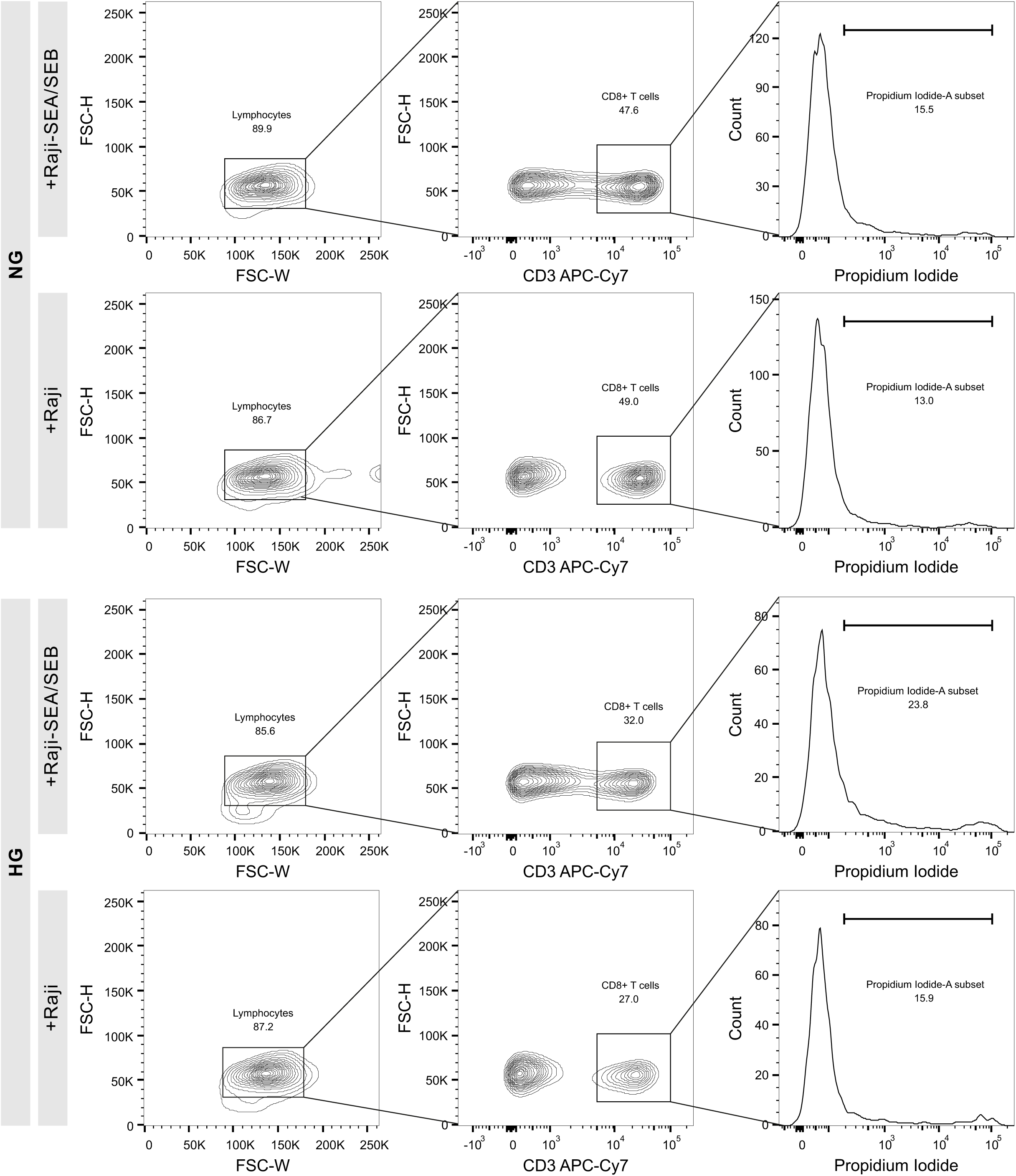
Gating strategy for CTL viability. Day 3 bead-stimulated primary human CD8+ T cells were used as CTLs. Raji cells were pulsed with or without SEA/SEB. CTLs and target cells were co-incubated for 16 hours at 37°C with 5% CO2. Then the cells were stained with APC/Cy7-αCD3 and BV-421-αCD19, followed by PI staining. Viable CTLs were gated as shown in the figure. One representative donor is shown out of 4 and 6 donors from 3 and 5 independent experiments for Raji and Raji-SEA/SEB, respectively.

## Notes

### Competing Interest Statement

The authors have declared no competing interest.

